# Water-related leaf traits associated with epiphytism in the Neotropical megadiverse genus *Anthurium* (Araceae)

**DOI:** 10.64898/2026.09.15.751687

**Authors:** Henriette F. Morgenroth, Gerhard Zotz, Nils Köster, H. Spencer Badet, Tiffany Knight, Jose B. Lanuza

## Abstract

**Background and Aims:** Epiphytes comprise approximately 10% of global vascular plant diversity and play key roles in many tropical ecosystems. They possess traits allowing survival under limited water availability (i.e. the epiphytic trait syndrome), but knowledge of these trait combinations is taxonomically biased and their generality remains unclear. The megadiverse genus *Anthurium* (Araceae), with ∼600 epiphytic species, offers an opportunity to address this gap by examining patterns of leaf trait covariation, their phylogenetic context, and differences among life forms.

**Methods:** A common garden experiment was conducted using 40 species from 13 sections spanning a gradient of epiphytic incidence. Twenty-one anatomical, morphological, and physiological leaf traits were quantified. Patterns of trait variation were assessed, and associations between leaf traits and epiphytism were evaluated both with and without accounting for phylogenetic relationships.

**Key Results:** Leaf trait variation was captured by two principal components. The first axis was negatively associated with stomatal density and size, leaf thickness, and leaf water content, while the second axis described a trade-off between leaf mass per area (LMA) and water loss rate. Epiphytic species generally exhibited smaller, more numerous stomata and higher LMA, consistent with adaptations for water retention. Epiphytism showed a phylogenetic signal; however, trait covariation patterns were not influenced by phylogenetic relatedness.

**Conclusions:** We found consistent associations between epiphytism and several water-related leaf traits in *Anthurium*, but some of these associations departed from general expectations of an epiphytic trait syndrome. Water-related traits showed variable phylogenetic signal and reflected multiple functional strategies for coping with intermittent water supply. These findings highlight the diversity of functional strategies underlying epiphytism and expand current understanding beyond well-studied taxa.

## Introduction

Vascular epiphytes, defined as non-parasitic plants that germinate and grow on other plants without contact with the soil (Benzing 1990; Zotz 2016) comprise approximately 10% of global vascular plant biodiversity. They are particularly prominent in neotropical regions where they locally constitute up to 50% of vascular species in humid rainforests (Kelly *et al*. 2004). Epiphytism occurs in 79 out of roughly 500 families of vascular plants, with Orchidaceae, Bromeliaceae and Polypodiaceae representing 86% of epiphytic diversity (Zotz, Weigelt, *et al*. 2021); but the families Ericaceae, Piperaceae, Gesneriaceae and Araceae are also key contributors (Zotz 2016). Despite the high diversity and broad taxonomic distribution of vascular epiphytes, the functional trait strategies associated with epiphytism remain biased towards a few dominant taxa (Hietz et al. 2021).

Epiphytes can play key ecological roles. The combined green biomass of vascular and non-vascular epiphytes can equal or even exceed that of tree leaves (Díaz *et al*. 2010) and they can play key roles in nutrient and water cycling (Gotsch *et al*. 2016). By modifying local temperature and humidity regimes (Stuntz, Simon, et al. 2002), they buffer microclimatic extremes and create habitats that are vital to many other taxa. For example, freshwater reservoirs in tank bromeliads provide breeding habitats for canopy-dwelling fauna like frogs or mosquitoes (Galindo-Leal *et al*. 2003; Quijano-Cuervo *et al*. 2021) or may host aquatic plants, such as carnivorous bladderworts (i.e., *Utricularia* spp., Zotz 2016). In addition, vascular epiphytes provide food and nesting material for birds (Gotsch *et al*. 2016) and insects (Kuanprasert *et al*. 1999; Ramírez 2019), and may engage in specialized ecological interactions (e.g., ant gardens; Davidson 1988). All in all, epiphyte abundance and diversity are often positively associated with those of other taxa (Stuntz, Ziegler, *et al*. 2002; Méndez-Castro *et al*. 2018), highlighting their central role in shaping forest ecosystem structure and function.

Growing in the canopy brings many advantages to plants, such as improved access to light, less herbivory, less competition, and less danger from flooding, trampling, and wildfires (Putz *et al*. 1995; Hietz *et al*. 2021). However, the lack of soil, the discontinuity of the surface they root upon, and the limited number of sites suitable for germination create difficulties for seed dispersal, seedling establishment and subsequent growth. Epiphytes often have limited access to essential nutrients, as well as reduced and highly variable water availability due to their reliance on atmospheric sources (Benzing 1990; Zotz 2016). To cope with these challenges, species exhibit a range of traits that enhance water and nutrient acquisition and retention in leaves, roots, and shoots. For instance, variation in leaf size (Zotz 2016), the presence of succulent leaves (Zhang *et al*. 2018) and low cuticular permeability (Helbsing *et al*. 2000) are common anatomical adaptations that limit water loss and/or improve water storage. Importantly, several studies have shown that seed dispersal and nutrient availability is not as limiting as water (Putz and Holbrook 1986; Zotz, Hietz, *et al*. 2021). Thus, traits in epiphytic plants are expected to be more tightly coupled to atmospheric water conditions compared to terrestrial plants (Mendieta-Leiva *et al*. 2020). Species composition and trait distributions are also likely to vary substantially across forest types, ranging from dry forests to wet cloud forests. Despite progress in understanding trait variation in epiphytes, especially in species-rich families like orchids and bromeliads, many taxonomic groups remain largely unexplored (Hietz *et al*. 2021). This highlights the need to evaluate whether the observed trait combinations and trade-offs can also be observed in these understudied taxonomic groups.

Aroids represent one of these understudied groups. Species in the family Araceae may exhibit multiple habits during ontogeny by developing adventitious roots and/or losing proximal shoot parts as they ascend host trees (Croat 1988; Zotz 2016). For instance, some species undergo ontogenetic shifts from an epiphytic to a terrestrial habit (Zotz et al. 2020). In addition, life forms can vary across habitats, highlighting that their ecological classification is context-dependent and needs to be considered across developmental stages. Consequently, substantial variability exists in the life forms assigned to species in the literature and in herbarium voucher descriptions. Although standardized nomenclature has been proposed to reduce this ambiguity (Zotz *et al*. 2020), including five forms of structurally dependent plants in the genus *Anthurium* (epiphytes, hemiepiphytes, nomadic vines, vines, and lithophytes), such discrete categories may not adequately capture the continuous nature of epiphytic dependence. Here, we move beyond categorical classification by incorporating a continuous measure of epiphytic incidence, which quantifies species’ position along the gradient from terrestrial to epiphytic growth (Zotz *et al*. 2023). This approach provides a more refined framework for linking ecological strategies to environmental conditions and, consequently, for detecting patterns in trait combinations and covariation.

This study examines to what extent epiphytic incidence is associated with water-related leaf traits in the neotropical genus *Anthurium*, building on earlier work that demonstrated intraspecific trait variation linked to life form in a species of this genus (Rada and Jaimez 1992). Among the roughly 4,600 species in the Araceae family (Boyce *et al*. 2025), the genus *Anthurium* comprises over 1,450 species (Plants of the World Online 2026). Excluding nomadic vines, there are around 700 species of Araceae that mostly grow as epiphytes, most of which belong to the genus *Anthurium* with c. 570 species (Reimuth and Zotz 2020; Zotz *et al*. 2020). The selection of traits is informed by previous studies (Holbrook and Putz 1996; Zhang *et al*. 2015; Hietz *et al*. 2021) and includes anatomical, morphological and physiological leaf traits associated with water retention. Because closely related species tend to share similar traits (de Bello *et al*. 2015), accounting for phylogenetic relationships can provide insights into the evolutionary processes shaping associations between life forms and functional traits. However, convergent evolution can also occur, in which unrelated species independently evolve similar traits in response to similar environmental pressures. Arising questions in this context are: (i) What are the major leaf trait covariation patterns in *Anthurium* concerning water relations? (ii) How is trait correlation influenced by evolutionary lineage? (iii) How do water-related leaf traits vary along the gradient of epiphytic incidence? Two hypotheses were specifically tested in this study: H1: water-related leaf traits vary systematically along the gradient of epiphytic incidence, reflecting increasing dependence on atmospheric water sources. H2: These trait-epiphytic incidence relationships are not phylogenetically conserved. By explicitly incorporating a continuous measure of epiphytic incidence, this study refines the evaluation of the proposed “epiphytic trait syndrome”, while acknowledging that such patterns may be context-dependent in epiphytic plants (Zotz 2016).

## Methods

### Study system

The study was conducted at the Botanic Garden Berlin from October to December of 2020. This common garden setting was chosen to minimize trait variation arising from environmental differences. A total of 21 water-related traits were studied in 40 species of the genus *Anthurium*, representing a wide range of life forms and phylogenetic groups (**Figure 1, Table S1**). These species represent 13 of the 19 botanical sections currently described for the genus (Carlsen et al., unpublished; Carlsen and Croat 2019; Croat and Carlsen 2020) and include taxa with terrestrial and epiphytic preference. Species were selected to capture the diversity of leaf traits and ecological strategies within the genus. All individuals were verified against morphological descriptions using taxonomic keys and sectional revisions (Madison 1978a; b; Croat 1991; Valadares and Sakuragui 2014), ensuring accurate species identification. Plants were grown under controlled greenhouse conditions (20 °C and 60% relative humidity) throughout the study period. Life forms were not treated as discrete categories but quantified using a continuous measure of epiphytic incidence (see life form section below). To preserve the integrity of the living collection, all measurements were designed to be minimally destructive while maximizing data acquisition.

**Figure 1.**
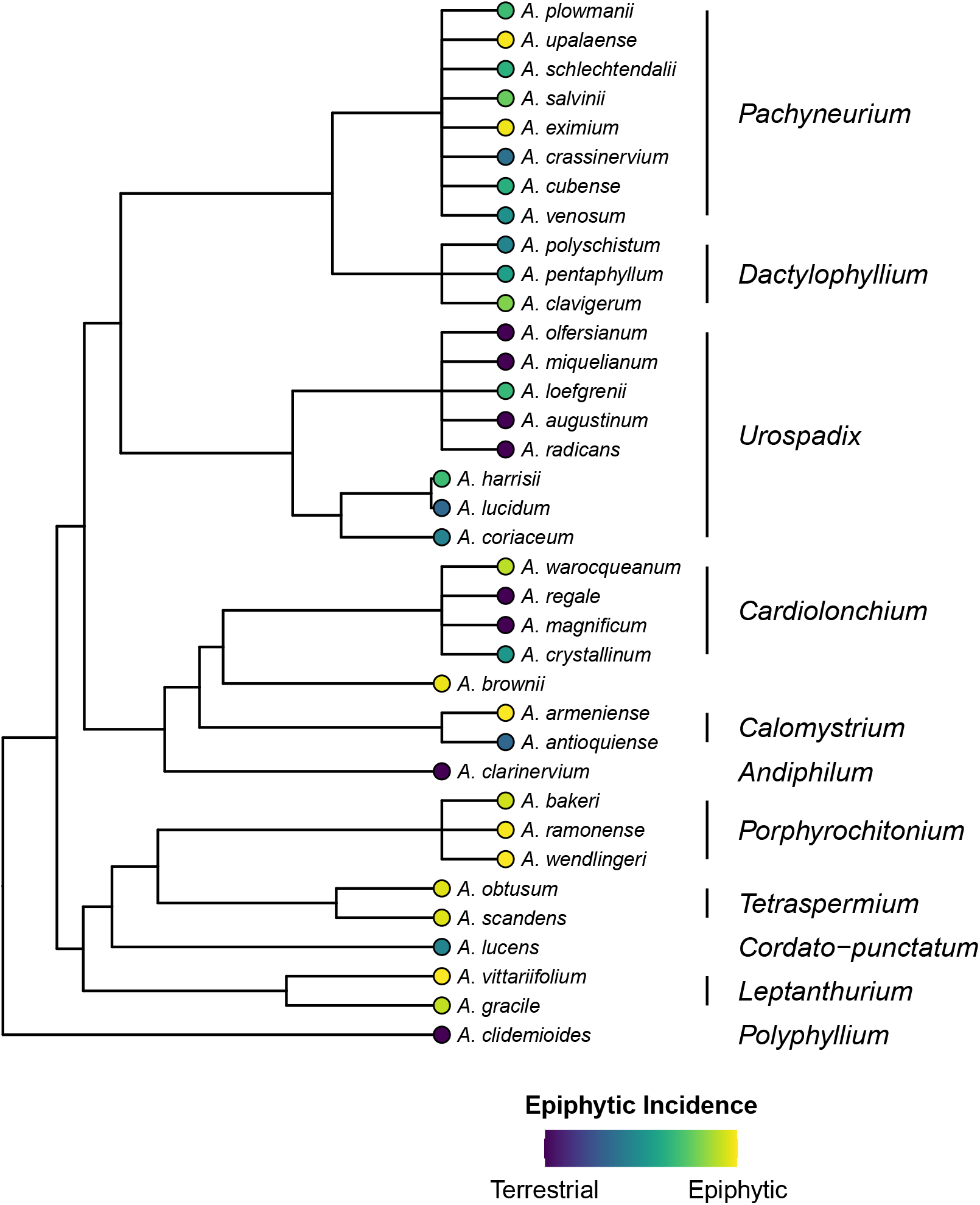
Phylogenetic tree of the *Anthurium* species used in this study. The different life forms are indicated with different colours at the tips of the branches. In addition, the most up-to-date *Anthurium* sections are shown on the right with vertical bars. Note that *Anthurium* brownii has not been placed into a section yet and the tree only contains 36 species of the 40 species used in this study due to the lack of phylogenetic information.

### Life forms

Life form classification in the Araceae remains challenging, as many species exhibit ontogenetic shifts and facultative changes in life form depending on environmental conditions (Zotz *et al*. 2020). Most existing classifications are categorical and do not capture this variability. To better capture the continuum of life form strategies, we quantified life form using a continuous measure of epiphytic incidence ranging from 0% (fully terrestrial) to 100% (fully epiphytic), based on literature data (Badet and Zotz, unpublished). Following the approach of Zotz *et al*. (2023), qualitative descriptions of species’ life forms were translated into quantitative estimates of epiphytic incidence (see **Table S2** for conversion criteria and **Table S3** for incidence level and sample sizes across species). This approach provides a more accurate description than previous analyses with strict categorical classifications that cannot capture the ecological diversity of life forms in this genus. To validate these estimates, life form was also assessed qualitatively using taxonomic resources, including specialist databases, herbarium vouchers, field images, and expert assessment (e.g. aroid.org, aroidpictures.fr, Botanic Garden Berlin collections, and field images provided by N.K.). The qualitative and semi-quantitative classification showed high similarity (83%), supporting the robustness of the approach. For all subsequent analyses, only the continuous epiphytic incidence values were used. Although the terms ‘life form’ and ‘growth form’ are inconsistently used in the literature, this work uses ‘life form’ throughout the whole text as the life form concept tries to put plant structure in an ecological context; in contrast to growth form, which generally describes the plant’s physiognomic habit (Ellenberg and Mueller-Dombois 1967; Zotz 2016).

### Leaf traits

In total, 21 water-related leaf traits were selected, representing anatomical, morphological and physiological dimensions (**Table 1**). Trait selection was based on previous studies examining functional trait variation in epiphytic plants (Holbrook and Putz 1996; Hietz and Briones 1998; Zhang *et al*. 2015; Hietz *et al*. 2021), and followed standardized protocols for plant trait measurement (Perez-Harguindeguy *et al*. 2016). The selected water-related traits were grouped into six categories: (i) water conservation traits, comprising anatomical features that reduce evaporative water loss (ii) water storage traits, which increase the capacity of leaves to retain water during periods of drought, (iii) water-use efficiency traits, reflecting the balance between carbon assimilation and water loss; (iv) traits related to water conservation and gas exchange, regulating transpiration while maintaining photosynthetic efficiency; (v) water transport traits, influencing hydraulic conductivity and thus the supply of water from uptake organs to the leaf; and (vi) water loss traits, representing unavoidable water loss through diffusion or transpiration.

**Table 1.**
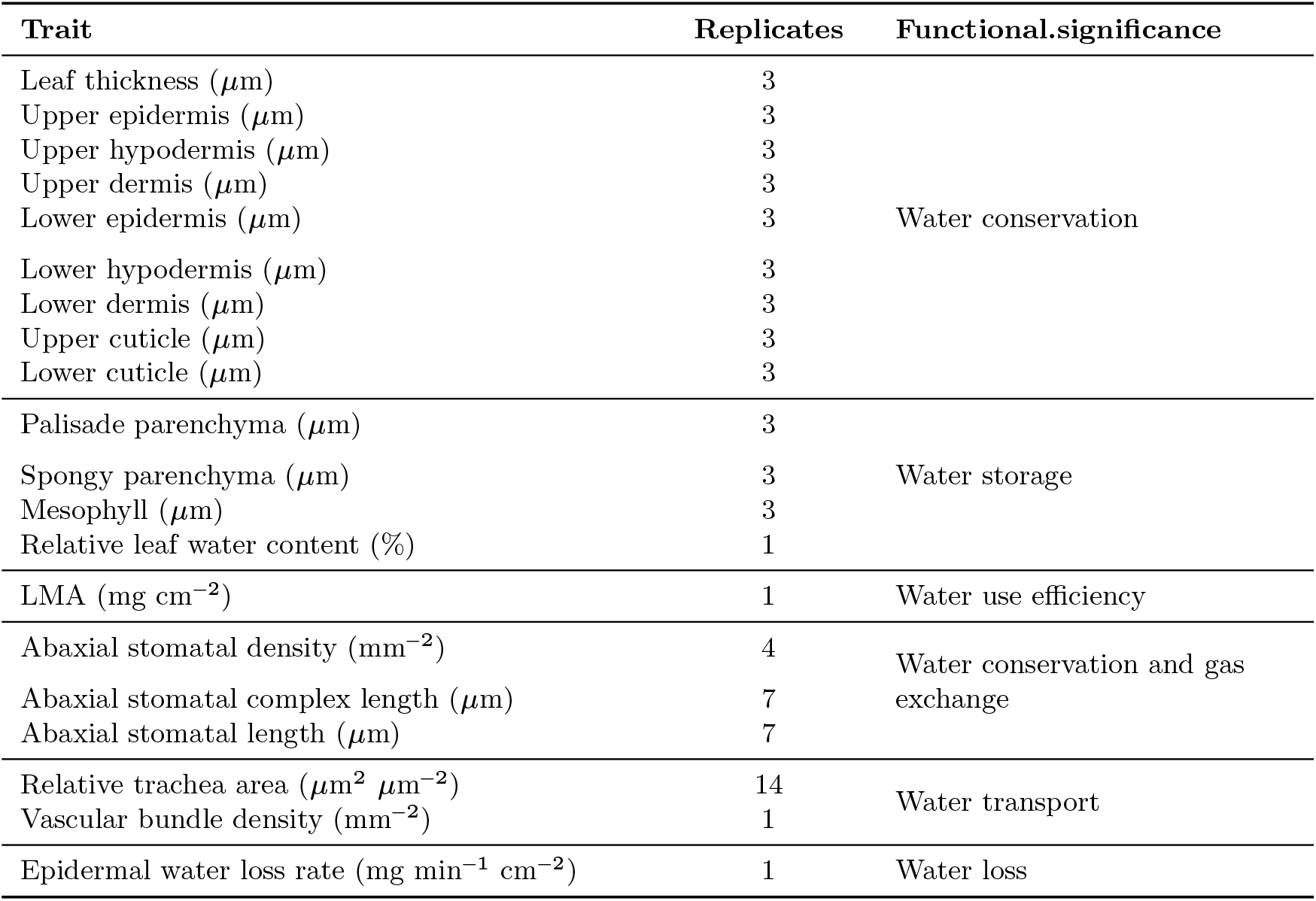
Water-related traits included in this study and their functional significance.

| Trait | Replicates | Functional.significance |
| --- | --- | --- |
| Leaf thickness ($\mu\text{m}$) | 3 | Water conservation |
| Upper epidermis ($\mu\text{m}$) | 3 | |
| Upper hypodermis ($\mu\text{m}$) | 3 | |
| Upper dermis ($\mu\text{m}$) | 3 | |
| Lower epidermis ($\mu\text{m}$) | 3 | |
| Lower hypodermis ($\mu\text{m}$) | 3 | |
| Lower dermis ($\mu\text{m}$) | 3 | |
| Upper cuticle ($\mu\text{m}$) | 3 | |
| Lower cuticle ($\mu\text{m}$) | 3 | |
| Palisade parenchyma ($\mu\text{m}$) | 3 | Water storage |
| Spongy parenchyma ($\mu\text{m}$) | 3 | |
| Mesophyll ($\mu\text{m}$) | 3 | |
| Relative leaf water content (%) | 1 |  |
| LMA ($\text{mg cm}^{-2}$) | 1 | Water use efficiency |
| Abaxial stomatal density ($\text{mm}^{-2}$) | 4 | Water conservation and gas exchange |
| Abaxial stomatal complex length ($\mu\text{m}$) | 7 | |
| Abaxial stomatal length ($\mu\text{m}$) | 7 | |
| Relative trachea area ($\mu\text{m}^2 \mu\text{m}^{-2}$) | 14 | Water transport |
| Vascular bundle density ($\text{mm}^{-2}$) | 1 | |
| Epidermal water loss rate ($\text{mg min}^{-1} \text{cm}^{-2}$) | 1 | Water loss |

Water conservation traits included leaf thickness and the thickness of epidermis, hypodermis, dermis and cuticle of the adaxial and abaxial surfaces of the leaf. Dermis was defined as the combination of the epidermis and hypodermis when both layers were present. Water storage traits included palisade parenchyma, spongy parenchyma, mesophyll (i.e. sum of palisade and spongy layers where both occurred), and leaf water content, calculated as the difference between saturated fresh mass (after overnight hydration) and dry mass (after drying at 80 °C for 4.5 days). Water use efficiency was represented by leaf mass per area (LMA). Traits related to water conservation and gas exchange included abaxial stomatal density (number of stomata per mm^2^), abaxial stomatal complex length (length of stomata including guard cells) and abaxial stomatal length (i.e. length of stomata excluding guard cells). Only abaxial measurements were considered, as only six of the 40 species were amphistomatic. Water transport traits included vascular bundle density (number of bundles per unit petiole area) and water transport capacity. The latter was estimated as total trachea area per petiole cross-sectional area, without distinguishing between tracheids and tracheae. Trachea area was calculated as the mean area of conduits within 14 vascular bundles (assuming elliptical shape based on major and minor diameters) and extrapolated to the total number of bundles per petiole. Water loss traits comprised stomatal and cuticular water loss rates, derived from drying curves under controlled conditions (23°C, 55% RH) monitored using HOBO data loggers (Onset Computer Corporation, Cape Cod, USA). Leaves were weighed at regular intervals over approximately two weeks, with higher temporal resolution during the initial drying phase (every 15 min during the first 120 min, followed by 2-4 measurements per day over the subsequent two weeks). Water loss rates were estimated as the slope of linear models where the response variable was area-normalized water loss as a function of time. This analysis was performed separately for two time intervals: the first 60 min, representing primarily stomatal water loss, and the period thereafter, representing predominantly cuticular (epidermal) water loss (Holbrook and Putz 1996). The initial 60 minutes likely captured stomatal closure for all species, as most studies report closing times from 30 to 40 minutes (Muchow and Sinclair 1989), but this may extend to 60 min or more in some cases (Khan *et al*. 2024). For comparability among species, relative water content (RWC) was calculated as: 

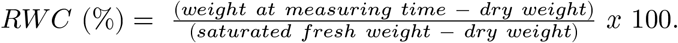

Anatomical traits of leaf blades were obtained by collecting leaf sections using a cork borer directly in the botanical garden. For petiole and leaf drying measurements (i.e. water transport and water loss traits, respectively), leaves were severed with a razor blade and immediately submerged in water, where final cuts were made to prevent embolism formation that could influence drying and wilting behavior. Leaves were then left with their petioles submerged in water overnight, and the cut surfaces were sealed the following day with Vaseline (< 0.1 g). Saturated fresh weight was determined using a digital scale (precision 0.01g, Sartorius Entris 2200-1S, Göttingen). Leaves were subsequently photographed alongside a reference scale to determine leaf area before being allowed to dry. All tissues (i.e. leaf blade and petiole) were processed in the lab where they were hand-cut and stained with Etzold’s simultaneous staining method (Fuchsia-Chrysoidine-Astra Blue) for one minute. Samples were then washed in 70% ethanol and stored overnight in distilled water. On the following day, tissues were photographed with a ZEISS Axiocam microscope under different magnifications depending on the size of the specific tissue. Images were stored in .TIFF format and analyzed using imageJ (Schneider *et al*. 2012). Stomatal traits were determined from epidermal impressions obtained by applying transparent nail polish to approximately 1 cm^2^ of both abaxial and adaxial leaf surfaces. Once dried, the impressions were mounted on slides and analyzed following the same imaging procedure.To minimize developmental variation, only mature plants of similar size were sampled, and measurements were conducted on the youngest fully expanded leaf of each individual.

### Phylogenetic tree

The phylogenetic tree was obtained using the R package GIFT (Denelle *et al*. 2023), which extracts species-level phylogenetic relationships from the global seed-plant megatree of Smith and Brown (2018). Because phylogenetic relationships within *Anthurium* remain incompletely resolved, the resulting topology was refined where possible using the molecular phylogenetic framework and most recent infrageneric classification of Carlsen and Croat (2019), complemented by ongoing phylogenetic work (M. Carlsen, personal comm., 6 November 2023). Species were placed according to these relationships when sufficient phylogenetic information was available; otherwise, uncertain relationships were retained as unresolved polytomies. Four species (*A. flavescens, A. willdenowii, A. sagittatum*, and *A. superbum*) for which no suitable taxonomic or phylogenetic information was available were excluded. The resulting tree was processed using ape (Paradis and Schliep 2019) and phytools (Revell 2024), and visualized using ggtree (Yu et al. 2017) and ggplot2 (Wickham 2011).

### Data analysis

Trait correlations were first used to identify and remove redundant variables with similar ecological relevance. Pairwise correlations were calculated using Spearman’s rank correlation coefficient, which is robust to outliers and does not assume normally distributed data. Patterns of trait covariation were explored using principal component analysis (PCA) to characterize the multivariate spectrum of water-related leaf traits. PCA was conducted using the function *PCA* in FactoMineR (Lê et al. 2008). Missing values (∼7% of the dataset) were imputed using mean substitution to retain all species in the analysis. To account for shared evolutionary history, a phylogenetically informed PCA was performed using *phyl*.*pca* (Revell 2024), incorporating Pagel’s to model phylogenetic signal. Results were qualitatively consistent with the non-phylogenetic PCA, indicating that phylogenetic relatedness does not strongly structure trait covariation (**Figure S1**). To assess the relationship between trait variation and epiphytic incidence, linear models were fitted with epiphytic incidence as the response variable and the first two principal components (PC1 and PC2) as predictors. In addition, associations between individual traits and epiphytic incidence were evaluated using Spearman’s rank correlation coefficient. All data processing and analyses were conducted in R version 4.2.2 (R Core Team 2022), primarily using tidyverse (Wickham et al. 2019).

## Results

### Life forms

Epiphytic incidence (%) varied considerably across species (mean = 57%; SD = 37). In total, 11 species were classified as primarily terrestrial (<33% epiphytic incidence), 10 as facultative epiphytes (between 33% and 66%), and 19 as epiphytes (>66%). Epiphytic incidence also varied across the distinct phylogenetic groups (**Figure 1**) and exhibited a moderate phylogenetic signal (= 0.58, P = 0.01), indicating that more closely related species are more likely to exhibit similar habit. For instance, in this dataset, section *Urospadix* contains mostly terrestrial species, section *Porphyrochitonium* contains only epiphytic species, whereas section *Pachyneurium* includes both epiphytic and facultatively epiphytic species.

### Leaf traits

The *Anthurium* species included in this study exhibited substantial interspecific trait variability (coefficient of variation, CV = 5% to 119%; **Table S4**). Traits with the lowest variability were relative leaf water content (CV = 5%), abaxial stomatal complex length (CV = 18%), and abaxial stomatal length (CV = 19%). In contrast, the most variable traits were stomatal water loss rate (CV = 119%), epidermal water loss rate (CV = 77%), and vascular bundle density (CV = 63%). This variation is reflected in the drying curves used to estimate water loss rates, which showed pronounced differences among species (**Figure S2**). Trait correlation analyses indicated that most trait pairs were weakly to moderately correlated (87%; values < |0.5|), while only 13% of trait pairs exhibited moderate to strong correlations (values |0.5|; **Figure 2**). Among these, only two trait pairs were negatively correlated, whereas all others showed positive associations.Ecologically redundant trait pairs with moderate to strong correlations included (i) upper and lower epidermis, (ii) upper and lower cuticle, (iii) leaf thickness and mesophyll, (iv) stomatal and epidermal water loss rates, and (v) stomatal and stomatal complex length. From each pair, upper epidermis, upper cuticle, leaf thickness, epidermal water loss rate, and stomatal length, respectively, were retained for further analyses based on ecological relevance and their contribution to overall trait variation. Some species exhibited distinct morphological and anatomical features, such as amphistomatic leaves (**Table S5**). As these traits occurred in only a small subset of species, they were not included in further statistical analyses.

**Figure 2.**
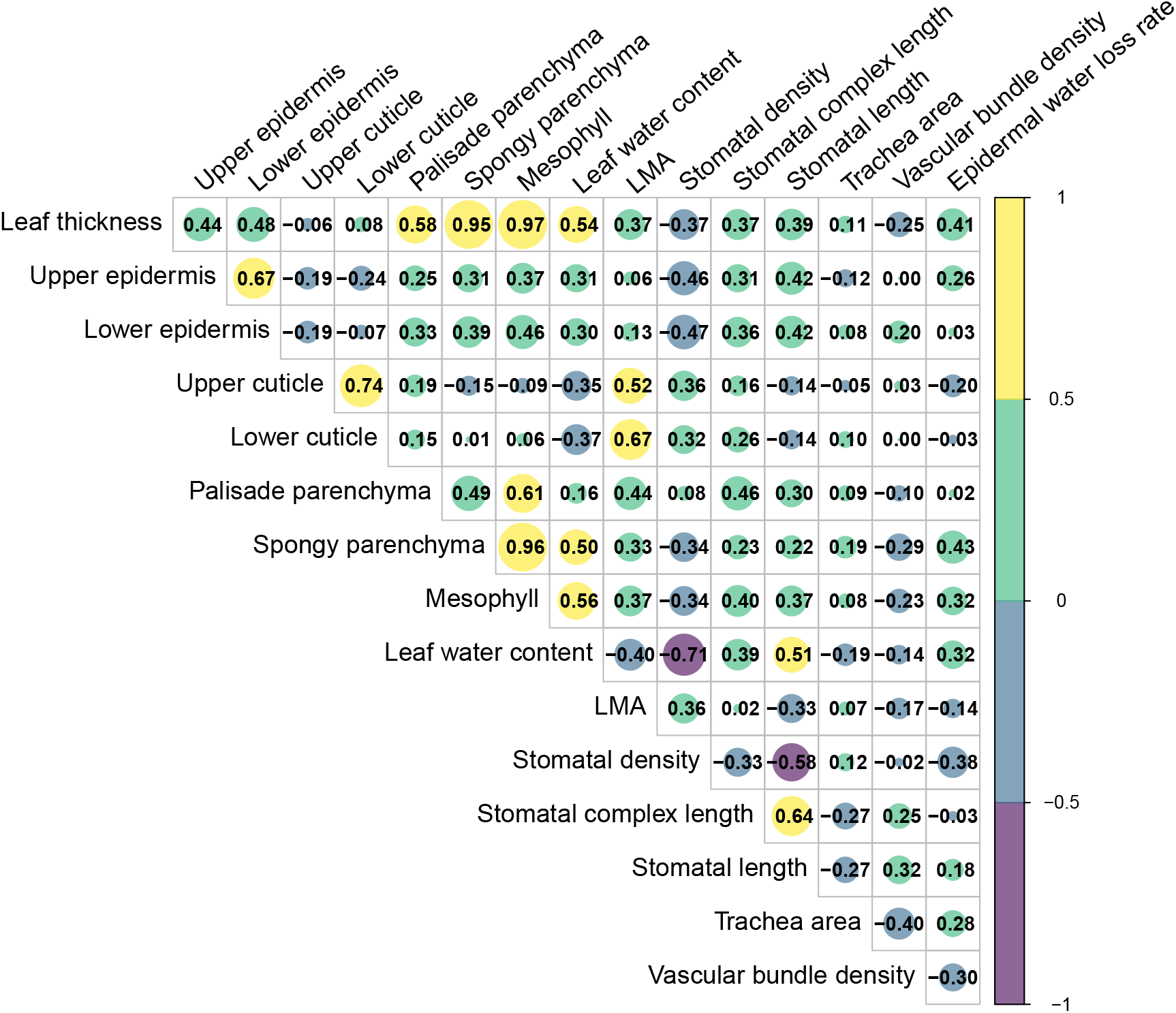
Correlogram of all traits considered in this study. Anatomical traits only occurring in a few species, i.e. with over 75% missing values, were excluded from the correlogram and further analyses. These were adaxial and abaxial hypodermis and dermis (sum of hypodermis and epidermis).

### Phylogenetic signal

Phylogenetic signal varied considerably across traits (**Table S6**). Traits exhibiting low phylogenetic signal (0.3) included upper epidermis thickness, palisade parenchyma thickness, trachea area, vascular bundle density, and stomatal water loss rate. Traits with moderate to high phylogenetic signal (> 0.3) included epiphytic incidence, leaf thickness, upper cuticle thickness, spongy parenchyma thickness, relative leaf water content, LMA, stomatal density, and stomatal length.

### Leaf traits and epiphytic incidence

Epiphytic incidence showed low to moderate correlations with relative leaf water-related traits (**Figure 3**). Only relative leaf water content, LMA, stomatal length, and stomatal water loss rate exhibited moderate correlations with epiphytic incidence (R 0.3). In general, species with higher epiphytic incidence tended to have lower relative leaf water content, higher LMA, smaller stomata, and lower stomatal water loss rates compared to species with lower epiphytic incidence.

**Figure 3.**
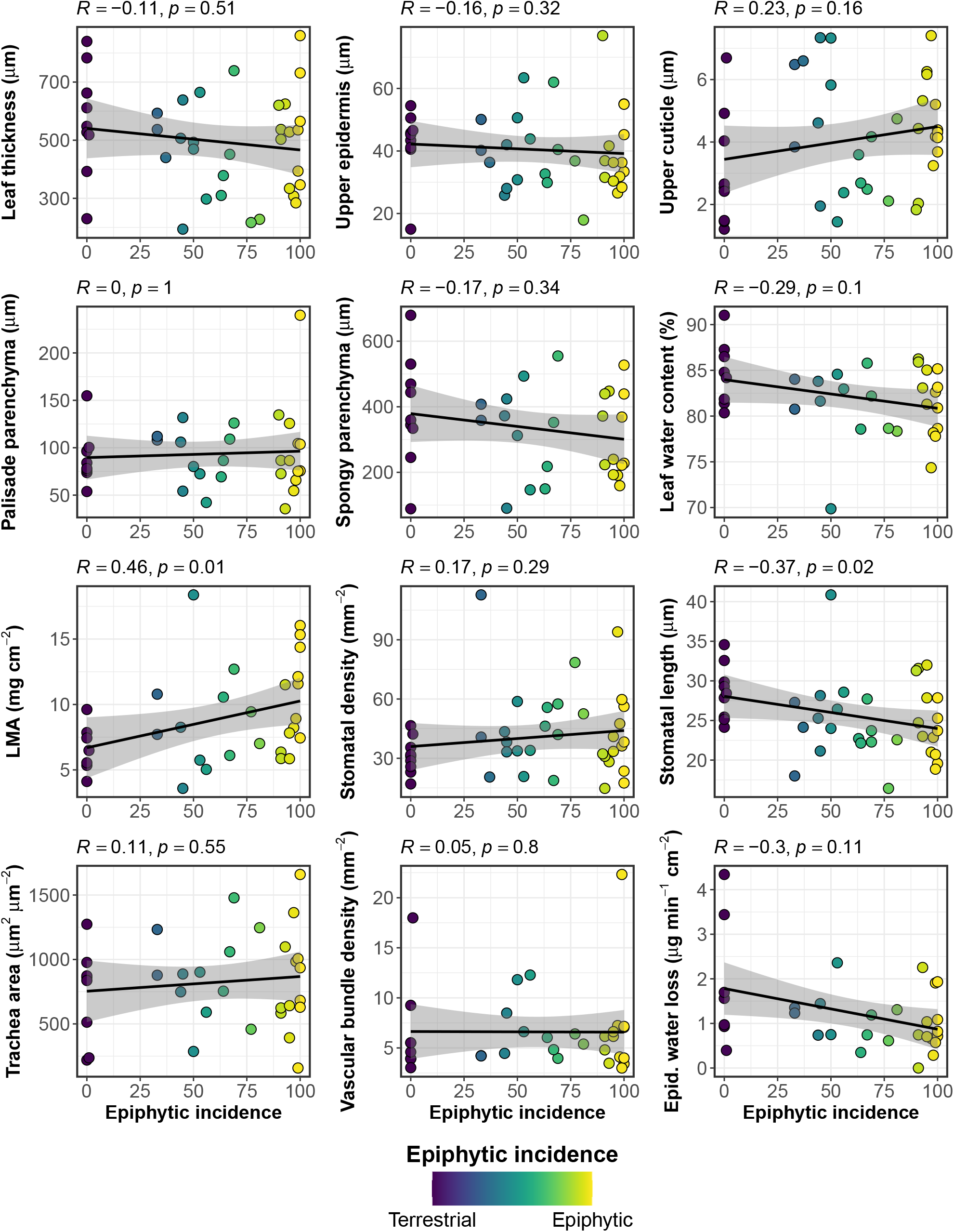
Different water-related leaf traits as a function of epiphytic incidence, the latter additionally illustrated by a colour gradient. On the x-axes, the degree of epiphytism is shown from 0% (terrestrial, blue) to 100% (holo-epiphytes, yellow).

#### Leaf trait spectrum

Approximately half of the total trait variation (47%) was captured by the first two principal components (PC1 and PC2; **Figure 4a**). The first axis explained 28% of the variation and showed a significant negative association with epiphytic incidence (= −6.56, *P* = 0.048; **Figure 4b**). The main traits contributing to this axis were leaf thickness, spongy parenchyma, relative leaf water content, stomatal density, and stomatal length (|loadings| > 2; **Table S7**). This axis reflects a trade-off between high stomatal density and thicker leaves with thicker parenchyma, larger stomata, and higher water content. For instance, *A. brownii* combined high stomatal density (94 mm ^2^) with relatively thin leaves (308 µm) and low water content (74%), whereas *A. clarinervium* showed a low stomatal density (17 mm ^2^) with thick leaves (662 µm) and high water content (100%).

**Figure 4.**
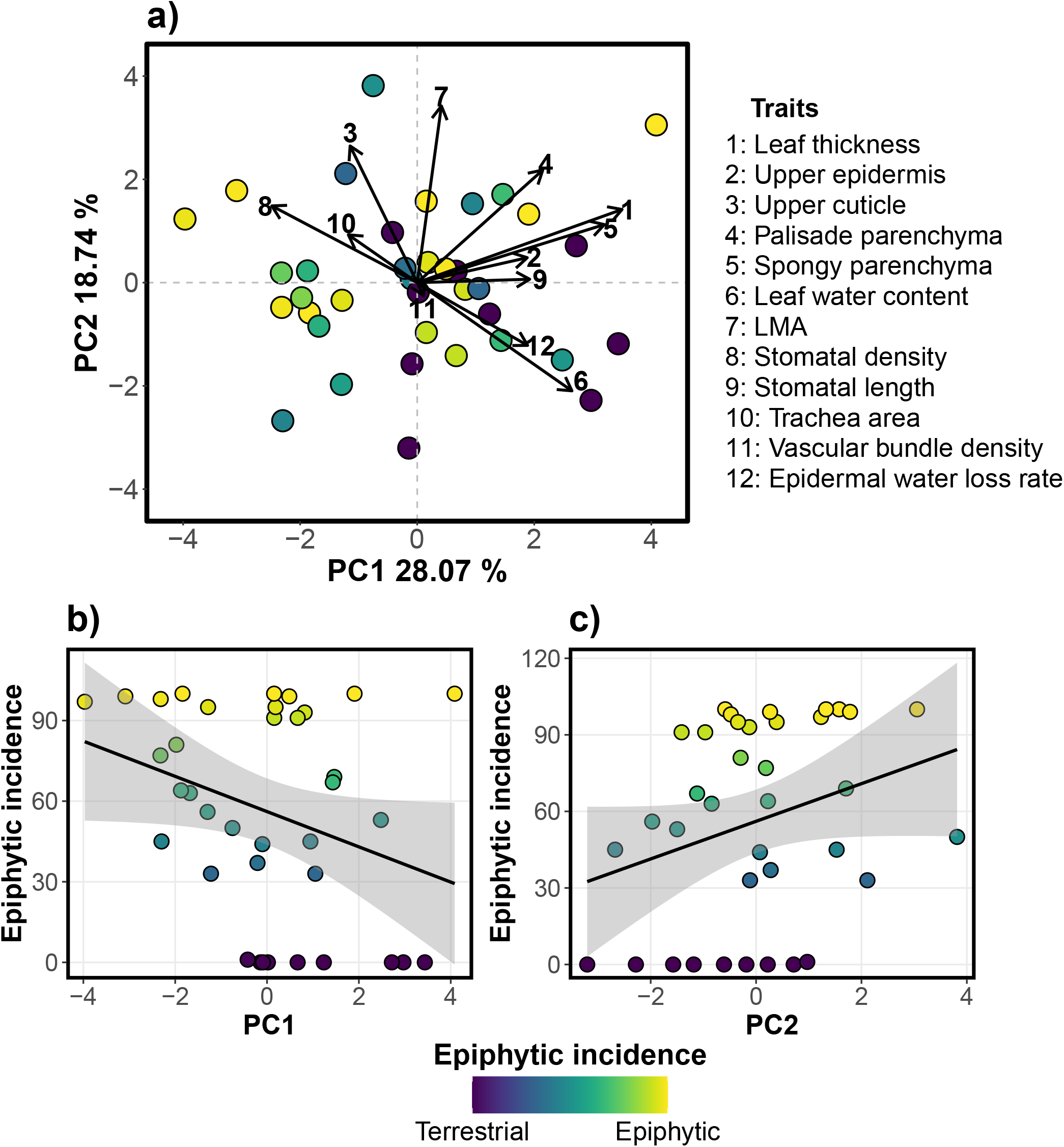
Water-related leaf trait variation in *Anthurium* species. Graph a) shows the principal component analysis (PCA) for a total of 12 traits with low functional redundancy and moderate to low correlation. Highly correlated traits were excluded to avoid overemphasizing their influence. The different points are coloured by the level of epiphytic incidence which ranges from 0% (terrestrial) to 100% (epiphytic). Graphs b) and c) show how epiphytic incidence changes across the main two axes of trait variation (PC1 and PC2, respectively).

The second axis explained 19% of the trait variation and showed a marginally positive association with epiphytic incidence (= 7.37, *P* = 0.068; **Figure 4c**). The main contributing traits were upper cuticle thickness, palisade parenchyma thickness, LMA, and stomatal water loss rate (|loadings| > 2; **Table S7**). Cuticle thickness, palisade parenchyma, and LMA were negatively associated with stomatal water loss rate. For example, *A. wendlingeri* exhibited a low water loss rate (2 × 10 mg min ^1^ cm ^2^) and high LMA (20 mg cm ^2^), partly driven by a thick cuticle (4 µm) and large palisade parenchyma (240 µm). In contrast, *A. clidemioides* showed a high water loss rate (3 × 10 mg min ^1^ cm ^2^), low LMA (4 mg cm ^2^), a thin cuticle (3 µm), and small palisade parenchyma (54 µm). A phylogenetically informed PCA yielded similar results (**Figure S1**; **Table S8**), indicating that evolutionary history did not strongly structure patterns of leaf trait covariation (0).

## Discussion

Water is a key limiting factor shaping the trait spectra of epiphytic plants (Hietz *et al*. 2021). However, its role in structuring life history strategies remains poorly understood across most epiphytic taxa. This study examines how variation in epiphytic incidence is associated with water-related leaf traits in the large but little-studied genus *Anthurium*. Using a common garden experiment with species that are characterized by different levels of epiphytic incidence and large interspecific trait variability, we show that (i) approximately half of the variation in water-related leaf traits is captured by two principal axes, (ii) these axes are largely independent of phylogenetic relatedness, and (iii) associations with epiphytic incidence are generally low to moderate. The first axis captures a trade-off between stomatal density and stomatal size, leaf thickness, and relative leaf water content, while the second axis reflects the trade-off between LMA and water loss rate. These trait associations vary along the gradient of epiphytic incidence, with species exhibiting higher epiphytic incidence tending to display trait combinations consistent with enhanced water retention, in line with patterns reported for other epiphytic taxa (Zhang *et al*. 2015; Zotz 2016). These results show that water-related leaf traits vary along the gradient of epiphytic incidence and are not strongly constrained by phylogeny, with associations being trait-specific and strongest for traits related to water loss.

The observed trade-off between stomatal density and size agrees with general findings in the plant kingdom (de Boer *et al*. 2016). Epiphytic *Anthurium* species tend to have smaller but more numerous stomata, which may reflect adaptations to changing hydraulic conditions by reducing transpiration rates or response times, preventing embolisms (Franks and Beerling 2009). Contrary to the results of the current study, most taxa with preference for epiphytic growth seem to have equally large or even larger but less dense stomata than terrestrial species (Campany *et al*. 2021; Hietz *et al*. 2021). Ferns and orchids appear to differ in the stomatal density-size trade-off, with dense but small stomata in ferns, and the opposite pattern in orchids (Hietz *et al*. 2021). Leaf thickness and relative leaf water content also varied along this stomatal size-density axis, with thicker leaves and higher relative leaf water content positively correlated with stomatal size, as also found by Gotsch *et al*. (2022) across various epiphytic taxa. While leaf thickness and especially relative leaf water content were generally lower in epiphytic plants, especially evident for leaf water content, previous research found the opposite trend (Zhang *et al*. 2015; Males and Griffiths 2017; Campany *et al*. 2021; Hietz *et al*. 2021).

This study also confirms the negative relationship between LMA and stomatal water loss rate (Wang *et al*. 2019). *Anthurium* species with higher epiphytic incidence showed higher LMA and reduced stomatal water loss rate. An increase in mass per area without changes in leaf thickness suggests higher leaf density (Poorter *et al*. 2009; De La Riva *et al*. 2016), likely due to densely packed cells, thicker cell walls, and/or increased sclerenchymatic tissue. While these structural traits may be costly (Witkowski and Lamont 1991), they can provide advantages for epiphytic growth by: (i) improving drought tolerance, as tissues are less prone to collapse under hydric stress (Tyree and Sperry 1989; Kramp *et al*. 2022); (ii) extending leaf lifespan (Ryser 1996), as they are more resistant to herbivory (Blumenthal *et al*. 2020) and structural damage (Alvarez-Clare and Kitajima 2007; Kitajima and Poorter 2010); and (iii) improving shade tolerance (Kitajima and Poorter 2010). However, low LMA values can also occur in epiphytes, as also reported for bromeliads (Males and Griffiths 2017), potentially reflecting differences in canopy position and associated light and water availability among species. Therefore, although the genus *Anthurium* generally follows existing trade-offs for epiphytes, their trait syndrome does not always correspond to general trends. These findings highlight the importance of incorporating understudied taxonomic groups to better capture the diversity of strategies of epiphytic species.

Using the most recent phylogenetic information available for the genus *Anthurium* (Carlsen and Croat 2019), we evaluated how the phylogenetic relationships between the different species influence epiphytic incidence and trait covariation patterns. Consistent with previous studies, epiphytism showed a significant phylogenetic signal (Watkins Jr and Cardelús 2012; Dubuisson *et al*. 2013; Frenzke *et al*. 2016; Males and Griffiths 2017). However, similar to the findings of Helsen *et al*. (2023) for ferns, trait covariation patterns remained largely unchanged when accounting for phylogenetic relationships. Although phylogenetic effects on trait covariation patterns were weak, several individual traits exhibited moderate to strong phylogenetic signal. Overall, only a subset of water-related leaf traits were statistically conserved, but phylogenetic values were generally high for most traits, highlighting the importance of evolutionary processes shaping leaf water-related traits in *Anthurium*. This result is supported by the seemingly recent radiation in distinct clades (Carlsen 2011) and their occurrence in different geographical regions (Reimuth and Zotz 2020), which likely reflect distinct speciation events influenced by unique ecological niches. While some taxonomic groups show strong phylogenetic conservatism in leaf traits of epiphytes (Helsen *et al*. 2023), this pattern is not consistent across taxa; for example, leaf traits in Ericaceae appear largely independent of phylogenetic structure (Khan *et al*. 2024). It is also worth noting that this study represents a limited subset of *Anthurium* diversity, and broader analyses incorporating genetic data may provide a more comprehensive understanding of trait variation across the genus.

Future research could expand trait coverage by including physiological traits such as transpiration rates in living plants (i.e., the time required to reach 70% relative leaf water content as a threshold for physiological damage; Zhang *et al*. 2018), and recovery capacity following desiccation (Zotz and Andrade 1998). Further investigation of these and other traits may clarify trait correlation patterns, such as the causes of higher leaf density in epiphytes (e.g. cell architecture and xylem structure; Guimarães *et al*. 2019; Pimenta *et al*. 2021). Exploring adaptations to other limiting factors, including light availability, nutrient levels, and structural support, may reveal a wide range of strategies related to photosynthetic capacity, chemical traits (Petter *et al*. 2016), and herbivore defense mechanisms (Guimarães *et al*. 2019). Beyond leaves, other organs remain understudied (Hietz *et al*. 2021), such as roots (water uptake; Werner et al. 2023; Tay et al. 2024), shoots (water storage and architecture (Göbel *et al*. 2020), and reproductive structures (Etl *et al*. 2017). Different life forms may also shape ecological interactions, though the extent to which they influence plant-plant or plant-animal interactions remains largely unexplored, including commensal plant associations (Pimenta *et al*. 2021) and plant-ant interactions (Croat 1988). Finally, examining variation across ontogenetic stages may help identify potential evolutionary bottlenecks (Ray 1987; Zotz 2000; Lorenzo *et al*. 2010).

The sample size in this study is adequate to detect broad ecological tendencies (Martínez-Abraín 2014). However, larger datasets could provide deeper insights into the epiphytic trait syndrome (Hulshof and Swenson 2010), intraindividual variability in large epiphytic plants (Herrera *et al*. 2015; Castro Sánchez-Bermejo *et al*. 2023), and intraspecific variation in facultative epiphytes, as well as the influence of natural habitats on trait adaptations (Rada and Jaimez 1992; Petter *et al*. 2016; Guzmán-Jacob *et al*. 2022). Increased sampling may also improve phylogenetic analyses (Werner *et al*. 2023; Tay *et al*. 2024), although trait associations can remain largely independent of phylogeny even at larger scales (Khan *et al*. 2024). A more comprehensive phylogenetic backbone would further strengthen trait-life form correlation analyses, as demonstrated by Werner *et al*. (2023). Expanding the taxonomic scope to related genera such as *Philodendron* and *Monstera* may also improve understanding of epiphytic strategies within Araceae (Croat 1988; Croat and Ortiz 2020). Most *Anthurium* species occur at relatively low heights on host trees, with only a few species reaching higher canopy positions (Croat 1988). Species such as *A. gracile* and members of section *Porphyrochitonium* occur higher in the canopy and possess traits associated with greater tolerance of exposed habitats, including lithophytic conditions. Although lithophytes were not examined here, these traits may allow occupation of various habitats with overlapping ecological requirements (Zotz and Einzmann 2023).

Our results generally confirm previously described trait correlation patterns (Hietz *et al*. 2021), placing species with higher epiphytic incidence toward the “resource retention under unfavorable conditions” end of the plant economic spectrum (Wright *et al*. 2004). This study provides a comprehensive analysis of water-related leaf traits in *Anthurium* and their relationship to plant life forms, highlighting distinct water-use strategies. Species with higher epiphytic incidence tend to have particular trait correlations, particularly in stomatal density and size, leaf thickness, and LMA, that support water retention. While these patterns align with expectations of the epiphytic trait syndrome, deviations from general trends in other taxa are also evident. Phylogenetic relationships influence some traits but do not strongly constrain trait expression in *Anthurium*. Future work should expand trait coverage and sampling to better understand intraspecific variability and ecological adaptation. Including related taxa such as *Philodendron* and *Monstera* may provide a broader perspective on epiphytic strategies within Araceae. Overall, this study contributes to a broader understanding of how functional traits and evolutionary processes interact to shape ecological strategies in epiphytic plants, while highlighting the value of using a continuous gradient of epiphytic incidence to investigate epiphytic trait variation.

## Acknowledgements

We thank Prof. Dr. Helge Bruelheide (Martin Luther University Halle-Wittenberg) for his support throughout this study. HM is also grateful to Dr. Helena Einzmann (Carl von Ossietzky Universität Oldenburg), Prof. Dr. Martin Röser (Martin Luther University Halle-Wittenberg), Maria Schauer, Prof. Dr. Julien Bachelier (Freie Universität Berlin), and Dr. Mónica M. Carlsen (Missouri Botanical Garden) for valuable discussions and assistance. Equally, we thank the horticultural staff of the Botanic Garden Berlin for maintaining the living collection of *Anthurium* as an invaluable resource for research. JBL was supported by a BioDivFund postdoctoral fellowship and a Juan de la Cierva fellowship (JDC2023-052760-I).

## References

Alvarez-Clare S, Kitajima K. 2007. Physical defence traits enhance seedling survival of neotropical tree species. Functional Ecology 21: 1044–1054.

de Bello F, Berg MP, Dias AT, et al. 2015. On the need for phylogenetic ‘corrections’ in functional trait-based approaches. Folia Geobotanica 50: 349–357.

Benzing DH. 1990. Vascular epiphytes: general biology and related biota. Cambridge University Press, Cambridge.

Blumenthal DM, Mueller KE, Kray JA, Ocheltree TW, Augustine DJ, Wilcox KR. 2020. Traits link drought resistance with herbivore defence and plant economics in semi-arid grasslands: The central roles of phenology and leaf dry matter content. Journal of Ecology 108: 2336–2351.

de Boer HJ, Price CA, Wagner-Cremer F, Dekker SC, Franks PJ, Veneklaas EJ. 2016. Optimal allocation of leaf epidermal area for gas exchange. New Phytologist 210: 1219–1228.

Boyce PC, Croat TB, Hay A. 2025. The Überlist of Araceae, Totals for Published and Estimated Number of Species in Aroid Genera. https://www.aroid.org/ueberlist.

Campany CE, Pittermann J, Baer A, et al. 2021. Leaf water relations in epiphytic ferns are driven by drought avoidance rather than tolerance mechanisms. Plant, Cell & Environment 44: 1741–1755.

Carlsen, M. M. (2011). Understanding the origin and rapid diversification of the genus Anthurium Schott (Araceae), integrating molecular phylogenetics, morphology and fossils. PhD dissertation, University of Missouri-St. Louis, St. Louis, Missouri, USA.

Carlsen MM, Croat TB. 2019. An analysis of the sectional classification of Anthurium (Araceae): comparing infrageneric groupings and their diagnostic morphology with a molecular phylogeny of the genus. Annals of the Missouri Botanical Garden 104: 69–82.

Castro Sánchez-Bermejo P, Davrinche A, Matesanz S, Harpole WS, Haider S. 2023. Within-individual leaf trait variation increases with phenotypic integration in a subtropical tree diversity experiment. New Phytologist 240: 1390–1404.

Croat TB. 1988. Ecology and life forms of Araceae. Aroideana 11: 4–55.

Croat TB. 1991. A revision of Anthurium section Pachyneurium (Araceae). Annals of the Missouri Botanical Garden 78: 539–855.

Croat TB, Carlsen MM. 2020. A new section of Anthurium: section Cordato-punctatum (Araceae), restricted to Central America. Novon 28: 46–50.

Croat TB, Ortiz OO. 2020. Distribution of Araceae and the diversity of life forms. Acta Societatis Botanicorum Poloniae 89: 1–23.

Davidson DW. 1988. Ecological studies of neotropical ant gardens. Ecology 69: 1138–1152.

De La Riva EG, Olmo M, Poorter H, Ubera JL, Villar R. 2016. Leaf mass per area (LMA) and its relationship with leaf structure and anatomy in 34 Mediterranean woody species along a water availability gradient. PloS one 11: e0148788.

Denelle P, Weigelt P, Kreft H. 2023. GIFT—An R package to access the Global Inventory of Floras and Traits. Methods in Ecology and Evolution 14: 2738–2748.

Díaz IA, Sieving KE, Pena-Foxon ME, Larraín J, Armesto JJ. 2010. Epiphyte diversity and biomass loads of canopy emergent trees in Chilean temperate rain forests: A neglected functional component. Forest Ecology and Management 259: 1490–1501.

Dubuisson J-Y, Bary S, Ebihara A, Carnero-Diaz E, Boucheron-Dubuisson E, Hennequin S. 2013. Epiphytism, anatomy and regressive evolution in trichomanoid filmy ferns (Hymenophyllaceae). Botanical Journal of the Linnean Society 173: 573–593.

Ellenberg, H. and D. Mueller-Dombois. 1967. A key to Raunkiaer plant life forms with revised subdivisions. Bericht über das Geobotanische Forschungsinstitut Rübel zu Zürich 37:56–73.

Etl F, Franschitz A, Aguiar AJ, Schönenberger J, Dötterl S. 2017. A perfume-collecting male oil bee? Evidences of a novel pollination system involving Anthurium acutifolium (Araceae) and Paratetrapedia chocoensis (Apidae, Tapinotaspidini). Flora 232: 7–15.

Franks PJ, Beerling DJ. 2009. Maximum leaf conductance driven by CO effects on stomatal size and density over geologic time. Proceedings of the National Academy of Sciences 106: 10343–10347.

Frenzke L, Goetghebeur P, Neinhuis C, Samain M-S, Wanke S. 2016. Evolution of epiphytism and fruit traits act unevenly on the diversification of the species-rich genus Peperomia (Piperaceae). Frontiers in Plant Science 7: 1145.

Galindo-Leal C, Cendo-Vazquez JR, Calderon R, Augustine J. 2003. Arboreal frogs, tank bromeliads and disturbed seasonal tropical forest. Contemporary Herpetology 1: 1–12.

Göbel, C. Y., B. O. Schlumpberger, and G. Zotz. 2020. What is a pseudobulb? – Towards a quantitative definition. International Journal of Plant Science 181:686–696.

Gotsch SG, Nadkarni N, Amici A. 2016. The functional roles of epiphytes and arboreal soils in tropical montane cloud forests. Journal of Tropical Ecology 32: 455–468.

Gotsch SG, Williams CB, Bicaba R, et al. 2022. Trade-offs between succulent and non-succulent epiphytes underlie variation in drought tolerance and avoidance. Oecologia 198: 645–661.

Guimarães AL de A, do Tanque PR, dos Santos Pyrrho A, de Macêdo Vieira AC. 2019. Toxicological and anatomical study of vegetative organs of Anthurium maricense Nadruz and Mayo (Araceae). Revista Agrogeoambiental 11: 87–106.

Guzmán-Jacob V, Guerrero-Ramírez NR, Craven D, et al. 2022. Broad-and small-scale environmental gradients drive variation in chemical, but not morphological, leaf traits of vascular epiphytes. Functional Ecology 36: 1858–1872.

Helbsing S, Riederer M, Zotz G. 2000. Cuticles of vascular epiphytes: efficient barriers for water loss after stomatal closure? Annals of Botany 86: 765–769.

Helsen K, Viana JL, Lin T-Y, Kuo L-Y, Zelený D. 2023. Functional-trait contrasts between terrestrial and epiphytic ferns in Taiwanese subtropical cloud forests. Journal of Vegetation Science 34: e13220.

Herrera CM, Medrano M, Bazaga P. 2015. Continuous within-plant variation as a source of intraspecific functional diversity: Patterns, magnitude, and genetic correlates of leaf variability in Helleborus foetidus (Ranunculaceae). American Journal of Botany 102: 225–232.

Hietz P, Briones O. 1998. Correlation between water relations and within-canopy distribution of epiphytic ferns in a Mexican cloud forest. Oecologia 114: 305–316.

Hietz P, Wagner K, Nunes Ramos F, et al. 2021. Putting vascular epiphytes on the traits map. Journal of Ecology 110: 340–358.

Holbrook N, Putz F. 1996. From epiphyte to tree: differences in leaf structure and leaf water relations associated with the transition in growth form in eight species of hemiepiphytes. Plant, Cell & Environment 19: 631–642.

Hulshof CM, Swenson NG. 2010. Variation in leaf functional trait values within and across individuals and species: an example from a Costa Rican dry forest. Functional ecology 24: 217–223.

Kelly DL, O’Donovan G, Feehan J, Murphy S, Drangeid SO, Marcano-Berti L. 2004. The epiphyte communities of a montane rain forest in the Andes of Venezuela: patterns in the distribution of the flora. Journal of Tropical Ecology 20: 643–666.

Khan G, Schepker H, Buhk N, Hahn C, Albach DC, Zotz G. 2024. Functional ecology and evolution of terrestrial and epiphytic species of Rhododendron section Schistanthe (Ericaceae). Perspectives in Plant Ecology, Evolution and Systematics 63: 125796.

Kitajima K, Poorter L. 2010. Tissue-level leaf toughness, but not lamina thickness, predicts sapling leaf lifespan and shade tolerance of tropical tree species. New Phytologist 186: 708–721.

Kramp RE, Liancourt P, Herberich MM, et al. 2022. Functional traits and their plasticity shift from tolerant to avoidant under extreme drought. Ecology 103: e3826.

Kuanprasert N, Kuehnle AR, Kuehnle A. 1999. Fragrance quality, emission, and inheritance in Anthurium species and hybrids. Aroideana 22: 48–62.

Lê S, Josse J, Husson F. 2008. FactoMineR: an R package for multivariate analysis. Journal of statistical software 25: 1–18.

Lorenzo N, Mantuano DG, Mantovani A. 2010. Comparative leaf ecophysiology and anatomy of seedlings, young and adult individuals of the epiphytic aroid Anthurium scandens (Aubl.) Engl. Environmental and Experimental Botany 68: 314–322.

Madison M. 1978a. The Anthurium leuconeurum confusion. Aroideana 1: 17–19.

Madison M. 1978b. The species of Anthurium with palmately divided leaves. Selbyana 2: 239–282.

Males J, Griffiths H. 2017. Functional types in the Bromeliaceae: relationships with drought-resistance traits and bioclimatic distributions. Functional Ecology 31: 1868–1880.

Martínez-Abraín A. 2014. Is the ‘n=30 rule of thumb’ of ecological field studies reliable? A call for greater attention to the variability in our data. Animal Biodiversity and Conservation 37: 95–100.

Méndez-Castro FE, Bader MY, Mendieta-Leiva G, Rao D. 2018. Islands in the trees: A biogeographic exploration of epiphyte-dwelling spiders. Journal of Biogeography 45: 2262–2271.

Mendieta-Leiva, G., P. Porada, and M. Y. Bader. 2020. Interactions of epiphytes with precipitation partitioning. Pages 133-146 in J. T. Van Stan II, E. Gutmann, and J. Friesen, editors. Precipitation partitioning by vegetation. Springer, Cham.

Muchow R, Sinclair T. 1989. Epidermal conductance, stomatal density and stomatal size among genotypes of Sorghum bicolor (L.) Moench. Plant, Cell & Environment 12: 425–431.

Paradis E, Schliep K. 2019. ape 5.0: an environment for modern phylogenetics and evolutionary analyses in R. Bioinformatics 35: 526–528.

Perez-Harguindeguy N, Diaz S, Garnier E, et al. 2016. Corrigendum to: New handbook for standardised measurement of plant functional traits worldwide. Australian Journal of Botany 64: 715–716.

Petter G, Wagner K, Wanek W, et al. 2016. Functional leaf traits of vascular epiphytes: vertical trends within the forest, intra-and interspecific trait variability, and taxonomic signals. Functional Ecology 30: 188–198.

Pimenta KM, Mayo S, Temponi LG, Mantovani A, Amorim AM. 2021. Anthurium bromelicola and A. sterilispadix (Araceae): two distinct bromeliad commensals with highly unusual inflorescence morphology endemic to Northeast Brazil. Plant Systematics and Evolution 307: 1–18.

Plants of the World Online. 2025. Plants of the World Online.

Poorter H, Niinemets Ü, Poorter L, Wright IJ, Villar R. 2009. Causes and consequences of variation in leaf mass per area (LMA): a meta-analysis. New Phytologist 182: 565–588.

Putz FE, Holbrook NM. 1986. Notes on the natural history of hemiepiphytes. Selbyana 9: 61–69.

Putz FE, Romano GB, Holbrook NM. 1995. Comparative phenology of epiphytic and tree-phase strangler figs in a Venezuelan palm savanna. Biotropica 27: 183–189.

Quijano-Cuervo LG, Méndez-Castro FE, Rao D, Escobar Sarria F, Negrete-Yankelevich S. 2021. Spatial relationships between spiders and their host vascular epiphytes within shade trees in a Mexican coffee plantation. Biotropica 53: 954–965.

R Core Team. 2022. R: A Language and Environment for Statistical Computing. Vienna, Austria: R Foundation for Statistical Computing.

Rada F, Jaimez R. 1992. Comparative ecophysiology and anatomy of terrestrial and epiphytic Anthurium bredemeyeri Schott in a tropical Andean cloud forest. Journal of Experimental Botany 43: 723–727.

Ramírez S. 2019. Pollinator specificity and seasonal patterns in the euglossine bee-orchid mutualism at La Gamba Biological Station. Acta ZooBot Austria 156: 171–181.

Ray TS. 1987. Cyclic heterophylly in Syngonium (Araceae). American Journal of Botany 74: 16–26.

Reimuth J, Zotz G. 2020. The biogeography of the megadiverse genus Anthurium (Araceae). Botanical Journal of the Linnean Society 194: 164–176.

Revell LJ. 2024. phytools 2.0: an updated R ecosystem for phylogenetic comparative methods (and other things). PeerJ 12: e16505.

Ryser P. 1996. The importance of tissue density for growth and life span of leaves and roots: a comparison of five ecologically contrasting grasses. Functional Ecology 10: 717–723.

Schneider CA, Rasband WS, Eliceiri KW. 2012. NIH Image to ImageJ: 25 years of image analysis. Nature methods 9: 671–675.

Smith SA, Brown JW. 2018. Constructing a broadly inclusive seed plant phylogeny. American Journal of Botany 105: 302–314.

Stuntz S, Simon U, Zotz G. 2002. Rainforest air-conditioning: The moderating influence of epiphytes on the microclimate in tropical tree crowns. International Journal of Biometeorology 46: 53–59.

Stuntz, S., C. Ziegler, U. Simon, and G. Zotz. 2002. Diversity and structure of the arthropod fauna within three canopy epiphyte species in Central Panama. Journal of Tropical Ecology 18:161–176.

Tay J, Werner J, Zotz G. 2024. Morphological diversity of the velamen radicum in the genus Anthurium (Araceae). Plant Biology 26: 679–690.

Taylor A, Zotz G, Weigelt P, et al. 2022. Vascular epiphytes contribute disproportionately to global centres of plant diversity. Global Ecology and Biogeography 31: 62–74.

Tyree MT, Sperry JS. 1989. Vulnerability of xylem to cavitation and embolism. Annual Review of Plant Physiology and Plant Molecular Biology 40: 19–36.

Valadares RT, Sakuragui CM. 2014. A new species of Anthurium (Araceae) sect. Urospadix subsect. Obscureviridia from Espírito Santo, eastern Brazil. Systematic Botany 39: 31–35.

Wang C, He J, Zhao T-H, et al. 2019. The smaller the leaf is, the faster the leaf water loses in a temperate forest. Frontiers in Plant Science 10: 58.

Watkins Jr J, Cardelús CL. 2012. Ferns in an angiosperm world: Cretaceous radiation into the epiphytic niche and diversification on the forest floor. International Journal of Plant Sciences 173: 695–710.

Werner JC, Albach DC, Can L, Zotz G. 2023. The velamen radicum is common in the genus Anthurium, both in the epiphytic and terrestrial species. Diversity 16: 18.

Wickham H. 2011. ggplot2. Wiley interdisciplinary reviews: computational statistics 3: 180–185.

Wickham H, Averick M, Bryan J, et al. 2019. Welcome to the Tidyverse. Journal of open source software 4: 1686.

Witkowski E, Lamont BB. 1991. Leaf specific mass confounds leaf density and thickness. Oecologia 88: 486–493.

Wright IJ, Reich PB, Westoby M, et al. 2004. The worldwide leaf economics spectrum. Nature 428: 821–827.

Yu G, Smith DK, Zhu H, Guan Y, Lam TT-Y. 2017. ggtree: an R package for visualization and annotation of phylogenetic trees with their covariates and other associated data. Methods in Ecology and Evolution 8: 28–36.

Zhang S-B, Dai Y, Hao G-Y, Li J-W, Fu X-W, Zhang J-L. 2015. Differentiation of water-related traits in terrestrial and epiphytic Cymbidium species. Frontiers in Plant Science 6: 260.

Zhang S-B, Yang Y, Li J, et al. 2018. Physiological diversity of orchids. Plant Diversity 40: 196–208.

Zotz G. 2000. Size-related intraspecific variability in physiological traits of vascular epiphytes and its importance for plant physiological ecology. Perspectives in Plant Ecology, Evolution and Systematics 3: 19–28.

Zotz G. 2016. Plants on Plants-the Biology of Vascular Epiphytes. Springer.

Zotz, G., JL Andrade. 1998. Water relations of two co-occurring epiphytic bromeliads. Journal of Plant Physiology 152:545–554.

Zotz G, Armenia L, Einzmann HJR. 2023. A new approach to an old problem: how to categorize the habit of ferns and lycophytes. Annals of Botany 132: 513–522.

Zotz G, Bautista-Bello A, Kohlstruck J, Weichgrebe T. 2020. Life forms in aroids natural variability vs. terminological confusion. Aroideana 43: 315–333.

Zotz G, Einzmann HJR. 2023. How epiphytic are filmy ferns? A semi-quantitative approach. Diversity 15: 270.

Zotz G, Hietz P, Einzmann HJR. 2021. Functional ecology of vascular epiphytes. Annual Plant Reviews Online 4: 869–906.

Zotz G, Weigelt P, Kessler M, Kreft H, Taylor A. 2021. EpiList 1.0: a global checklist of vascular epiphytes. Ecology 102: e03326.

