## Supplementary Material for "Water-related leaf traits associated with epiphytism in the Neotropical megadiverse genus *Anthurium* (Araceae)"

**Table S1.** List of species used in this study, including the most up-to-date species names (according to World Flora Online).

**Species**

*Anthurium antioquiense* Engl.

*Anthurium armeniense* Croat

*Anthurium augustinum* Koch & Lauche

*Anthurium bakeri* Hook. f.

*Anthurium brownii* Mast.

*Anthurium clarinervium* Matuda

*Anthurium clavigerum* Poepp.

*Anthurium clidemioides* Standl.

*Anthurium coriaceum* G. Don

*Anthurium crassinervium* (Jacq.) Schott

*Anthurium crystallinum* Linden & André

*Anthurium cubense* Engl.

*Anthurium eximium* Engl.

*Anthurium flavescens* Poepp.

*Anthurium gracile* (Rudge) Lindl.

*Anthurium harrisii* (Graham) G. Don

*Anthurium loefgrenii* Engl.

*Anthurium lucens* Standl.

*Anthurium lucidum* Kunth

*Anthurium magnificum* Linden

*Anthurium miquelianum* K. Koch & Augustin

*Anthurium obtusum* (Engl.) Grayum subsp. obtusum

*Anthurium olfersianum* Kunth

*Anthurium pentaphyllum* (Aubl.) G. Don

*Anthurium plowmanii* Croat

*Anthurium polyschistum* R. E. Schult. & Idrobo

*Anthurium radicans* K. Koch & Haage

*Anthurium ramonense* K. Krause

*Anthurium regale* Linden

*Anthurium sagittatum* (Sims) G. Don

*Anthurium salvinii* Hemsl.

*Anthurium scandens* (Aubl.) Engl. subsp. scandens

*Anthurium schlechtendalii* Kunth

*Anthurium superbum* Madison

*Anthurium upalaense* Croat & R. A. Baker

*Anthurium venosum* Griseb.

*Anthurium vittariifolium* Engl.

*Anthurium warocqueanum* T. Moore

*Anthurium wendlingeri* G. M. Barroso

*Anthurium willdenowii* Kunth

**Table S2**. Verbal key used by Badet and Zotz (unpublished), similar to Zotz et al. (2023), to assign quantitative values of epiphytic incidence.

| **Verbal information** | **Assigned epiphytic incidence** |
| --- | --- |
| epiphyte and lithophyte,  epiphyte or terrestrial,  facultative epiphyte (50% to T),  terrestrial to epiphytic | 50% |
| epiphyte, lithophyte, or  terrestrial | 33% |
| holoepiphyte, epiphyte | 100% |
| obligatory epiphyte | 100% |
| rarely, seldom epiphyte | 5% |
| very rarely, accidental epiphyte,  exceptionally | 1% |
| also as, sometimes, occasionally,  uncommonly | 15% |
| commonly, generally, mainly,  mostly, preferentially, primarily,  usually, predominantly, typically | 85% |
| often, frequently, most  frequently, more commonly | 67% |
| less frequently, less commonly | 33% |

**Table S3.** Sample size (number of literature sources with information on habit) and average epiphytic incidence from Badet and Zotz (unpublished) for the 40 species included in this study. See Zotz et al. (2023) and Table S1 for methodological details.

| **Species** | **Sample size** | **Epiphytic incidence** |
| --- | --- | --- |
| *A. antioquiense* | 3 | 33 |
| *A. armeniense* | 3 | 100 |
| *A. augustinum* | 1 | 0 |
| *A. bakeri* | 11 | 94 |
| *A. brownii* | 11 | 97 |
| *A. clarinervium* | 3 | 0 |
| *A. clavigerum* | 17 | 85 |
| *A. clidemioides* | 9 | 78 |
| *A. coriaceum* | 5 | 44 |
| *A. crassinervium* | 9 | 59 |
| *A. crystallinum* | 7 | 60 |
| *A. cubense* | 11 | 62 |
| *A. eximium* | 5 | 98 |
| *A. flavescens* | 3 | 91 |
| *A. gracile* | 31 | 93 |
| *A. harrisii* | 13 | 70 |
| *A. loefgrenii* | 2 | 0 |
| *A. lucens* | 10 | 45 |
| *A. lucidum* | 1 | 33 |
| *A. magnificum* | 2 | 0 |
| *A. miquelianum* | 2 | 0 |
| *A. obtusum* | 16 | 96 |
| *A. olfersianum* | 1 | 0 |
| *A. pentaphyllum* | 31 | 60 |
| *A. plowmanii* | 4 | 69 |
| *A. polyschistum* | 4 | 34 |
| *A. radicans* | 1 | 0 |
| *A. ramonense* | 13 | 100 |
| *A. regale* | 4 | 0 |
| *A. sagittatum* | 3 | 0 |
| *A. salvinii* | 13 | 80 |
| *A. scandens* | 61 | 93 |
| *A. schlechtendalii* | 16 | 64 |
| *A. superbum* | 6 | 92 |
| *A. upalaense* | 9 | 99 |
| *A. venosum* | 2 | 50 |
| *A. vittariifolium* | 4 | 100 |
| *A. warocqueanum* | 2 | 90 |
| *A. wendlingeri* | 10 | 100 |
| *A. willdenowii* | 2 | 50 |

**Table S4.** Summary of the main water-related leaf traits across all species, including units, numerical range (minimum-maximum), mean values, and coefficients of variation (CV).

| **Traits** | **Units** | **Numerical range** | **Mean** | **CV** |
| --- | --- | --- | --- | --- |
| Leaf thickness | μm | 194-860 | 498 | 35 |
| Upper epidermis | μm | 15-77 | 40 | 31 |
| Lower epidermis | μm | 16-70 | 33 | 38 |
| Upper cuticle | μm | 1-7 | 4 | 46 |
| Lower cuticle | μm | 1-7 | 3 | 53 |
| Palisade parenchyma | μm | 36-240 | 93 | 40 |
| Spongy parenchyma | μm | 89-679 | 336 | 43 |
| Mesophyll | μm | 0-775 | 420 | 42 |
| Leaf water content | % | 0.7-0.9 | 0.8 | 5 |
| LMA | mg cm^−2^ | 4-18 | 9 | 41 |
| Abaxial stomatal density | mm^−2^ | 15-113 | 41 | 50 |
| Abaxial stomatal complex length | μm | 34-79 | 49 | 18 |
| Abaxial stomatal length | μm | 16-41 | 26 | 19 |
| Trachea area | μm² μm^−2^ | 158-1659 | 820 | 45 |
| Vascular bundle density | mm^−2^ | 3-22 | 7 | 63 |
| Epidermal water loss rate | mg min^−1^ cm^−2^ | 0-0.01 | 5 | 74 |

**Table S5.** Species exhibiting rare morphological and anatomical features. These traits occurred only in a small subset of species and were therefore excluded from the statistical analyses. They are reported here as a starting point for future detailed investigations.

| **Species** | **Amphistomatic** | **Undiff. Mesophyll** | **Hypodermis** | **Petiole Absent Sheath** | **Bilateral Palisade Parenchyma** | **Epiderm. Cells Papillose** | **Ruffled Leaves** |
| --- | --- | --- | --- | --- | --- | --- | --- |
| *A. augustinum* |  |  |  |  |  | X |  |
| *A. clarinervium* | X |  |  |  |  | X |  |
| *A. clidemioides* |  |  |  |  |  | X | X |
| *A. coriaceum* | X |  |  |  |  |  |  |
| *A. crassinervium* |  | X | X |  |  |  |  |
| *A. crystallinum* | X |  |  |  |  | X |  |
| *A. cubense* |  | X | X |  |  |  |  |
| *A. flavescens* |  |  |  | X |  |  |  |
| *A. gracile* |  | X | X |  |  |  |  |
| *A. magnificum* |  |  |  |  |  | X |  |
| *A. obtusum* |  | X | X | X |  |  |  |
| *A. pentaphyllum* |  |  |  | X |  |  |  |
| *A. polyschistum* |  |  |  |  |  |  | X |
| *A. radicans* |  |  |  | X |  |  |  |
| *A. salvinii* |  | X | X |  |  |  |  |
| *A. scandens* |  |  |  | X |  |  |  |
| *A. superbum* |  | X | X |  |  |  |  |
| *A. upalaense* |  | X | X |  |  |  |  |
| *A. venosum* |  | X | X |  |  |  |  |
| *A. vittariifolium* | X | X | X |  |  |  |  |
| *A. warocqueanum* |  |  |  |  |  | X |  |
| *A. wendlingeri* | X |  |  |  | X |  |  |

**Table S6.** Phylogenetic signal of epiphytic incidence and water-related leaf traits. Asterisks indicate significance levels (*P* < 0.05, P < 0.01).

| **Traits** | **Lambda** | **P-value** |
| --- | --- | --- |
| Epiphytic incidence | 0.58 | <0.05* |
| Leaf thickness | 0.61 | <0.01** |
| Upper epidermis | 0.26 | 0.15 |
| Upper cuticle | 0.72 | 0.07 |
| Palisade parenchyma | <0.01 | 1.00 |
| Spongy parenchyma | 0.62 | <0.05* |
| Leaf water content | 0.88 | <0.05* |
| LMA | 0.64 | 0.22 |
| Stomatal density | 0.70 | 0.09 |
| Stomatal length | 0.78 | <0.01** |
| Trachea area | 0.22 | 0.65 |
| Vascular bundle density | 0.09 | 0.61 |
| Epidermal water loss rate | <0.01 | 1.00 |

**Table S7.** Loadings of the two main axes of trait variation (PC1 and PC2) for the main uncorrelated water-related leaf traits.

| **Traits** | **PC1** | **PC2** |
| --- | --- | --- |
| Leaf thickness | 3.50 | 1.42 |
| Upper epidermis | 1.88 | 0.48 |
| Upper cuticle | -1.15 | 2.65 |
| Palisade parenchyma | 2.14 | 2.18 |
| Spongy parenchyma | 3.20 | 1.12 |
| Leaf water content | 2.64 | -2.10 |
| LMA | 0.41 | 3.42 |
| Stomatal density | -2.48 | 1.49 |
| Stomatal length | 1.92 | 0.07 |
| Trachea area | 0.11 | -0.25 |
| Vascular bundle density | -1.18 | 0.93 |
| Epidermal water loss rate | 1.89 | -1.22 |

**Table S8.** Loadings of the two main axes of trait variation (PC1 and PC2) from the phylogenetically informed principal component analysis for the main uncorrelated water-related leaf traits.

| **Traits** | **PC1** | **PC2** |
| --- | --- | --- |
| Leaf thickness | 3.33 | -1.93 |
| Upper epidermis | 2.39 | 0.13 |
| Upper cuticle | -1.40 | -2.30 |
| Palisade parenchyma | 2.36 | -2.27 |
| Spongy parenchyma | 3.07 | -1.80 |
| Leaf water content | 3.38 | 1.00 |
| LMA | 0.34 | -3.33 |
| Stomatal density | -2.64 | -1.59 |
| Stomatal length | 3.02 | 1.35 |
| Trachea area | -1.15 | -1.47 |
| Vascular bundle density | -0.69 | 0.88 |
| Stomatal water loss rate | 1.65 | 2.19 |

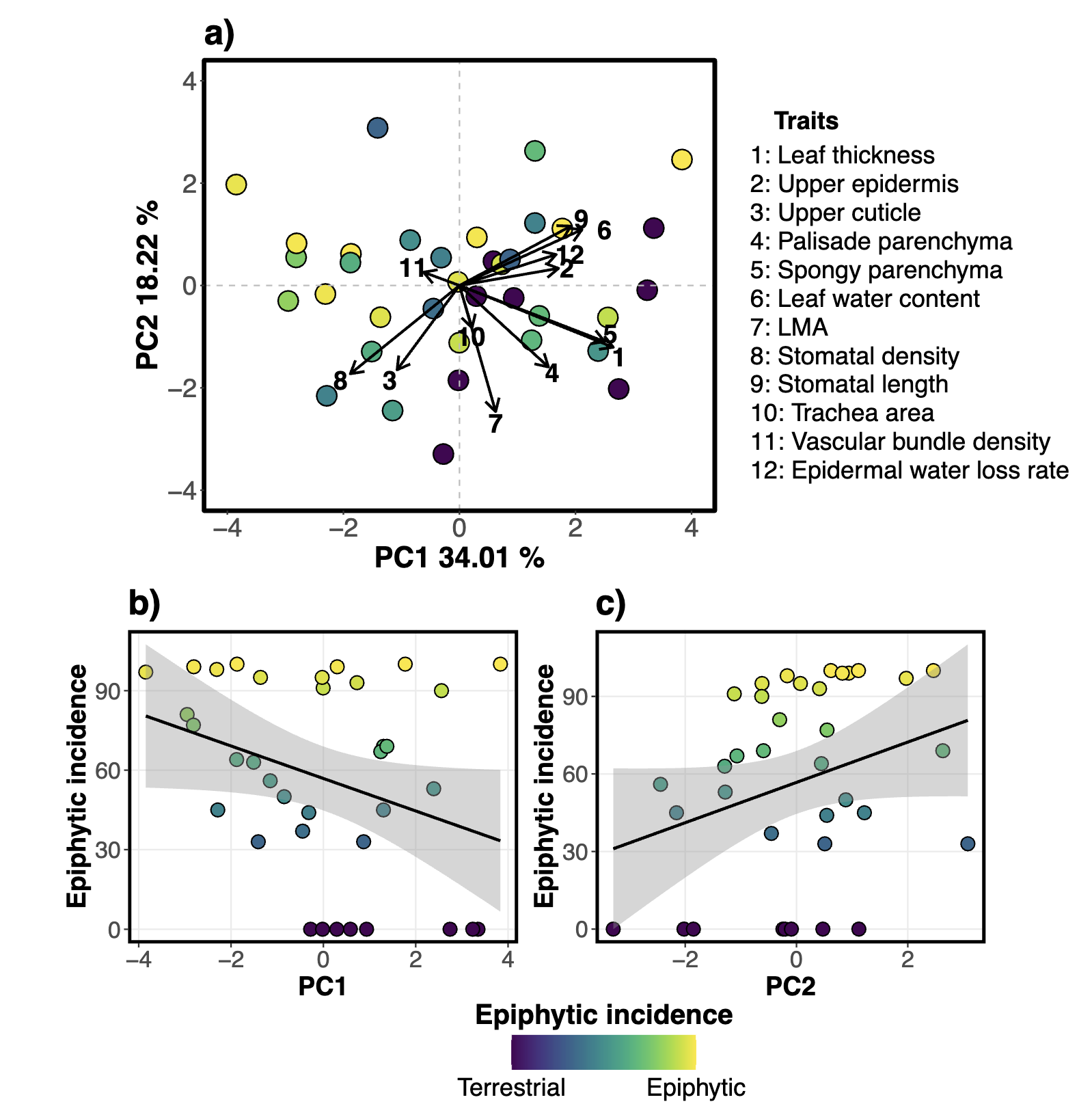

**Figure S1.** Phylogenetically corrected water-related leaf trait variation in *Anthurium* species. Graph a) shows the phylogenetic principal component analysis (pPCA) for a total of 12 traits with low functional redundancy and moderate to low correlation. Highly correlated traits were excluded to avoid overemphasizing their influence. The different points are coloured by the level of epiphytic incidence, which ranges from 0% (terrestrial) to 100% (epiphytic). Panels (b) and (c) show the relationship between epiphytic incidence and the first two axes of trait variation (PC1 and PC2, respectively). Black lines represent fitted linear regressions, and shaded areas indicate 95% confidence intervals.

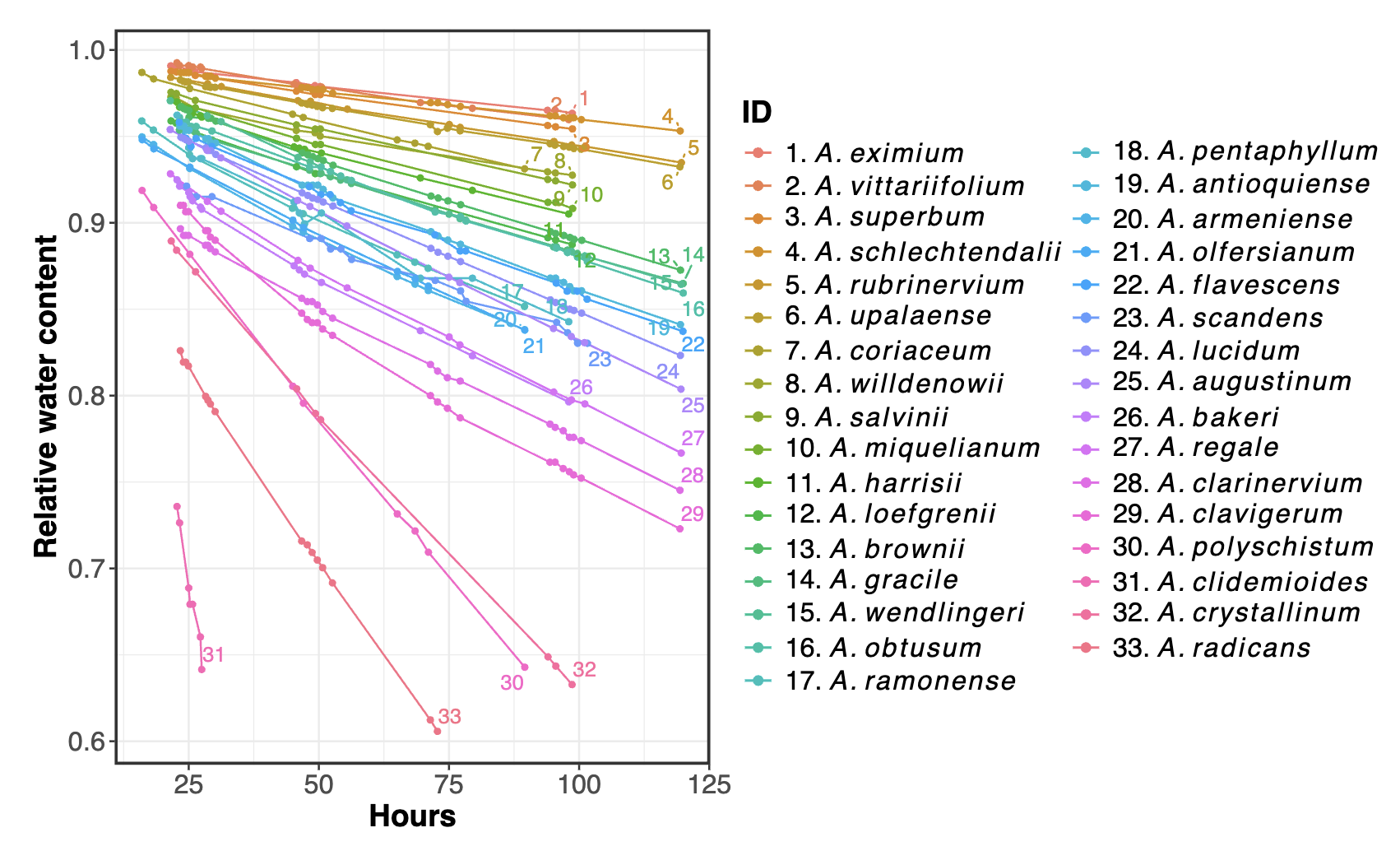

**Figure S2.** Leaf drying curves (water loss) for 33 out of 40 *Anthurium* species. Relative water loss over time is displayed, starting at 15 hours after cutting, when stomata are considered fully closed, and continuing until five days after cutting. One hundred percent corresponds to the full water content (saturated fresh weight) of the leaves. Curves are capped when relative water content (RWC) falls below 60%, as leaves are considered dead at this point. All samples were dried under constant environmental conditions.
